# Evaluating sequence-to-function deep learning models for ancestry-stratified regulatory variant effect prediction using multi-ancestry blood eQTLs

**DOI:** 10.64898/2026.06.22.730889

**Authors:** Xinyu Sun, Makaela Mews, Nicholas R. Wheeler, Penelope Benchek, Tianjie Gu, Lissette Gomez, Yousef Mustafa, Li-San Wang, Yuk Yee Leung, Gerard D. Schellenberg, Margaret A. Pericak-Vance, Jonathan L. Haines, Anthony J. Griswold, William S. Bush

## Abstract

**Background:** Sequence-to-function (S2F) deep learning models are increasingly used to prioritize non-coding regulatory variants, but their behavior across ancestrally diverse populations remains unclear. Because both training data and reference resources are heavily European-centered, multi-ancestry benchmarks are needed to determine whether S2F scores capture regulatory effects consistently across populations with different allele-frequency and LD patterns.

**Methods:** We evaluated Borzoi and AlphaGenome using whole blood eQTL data from the MAGENTA cohort, including African American (AA; *N* = 224), Caribbean Hispanic (CH; *N* = 209), and Non-Hispanic White (NHW; *N* = 235) participants. Model predictions were benchmarked against sampled nominal eQTLs and ancestry-stratified SuSiE fine-mapped variants using Spearman correlation, direction concordance, inter-model convergence, and distance-matched AUROC, with sensitivity analyses for minor allele frequency and comparison-set definition. We also compared FILER functional annotation overlap among high-Posterior Inclusion Probability (PIP) variants across ancestries.

**Results:** Both models showed weak agreement with nominal eQTL effect sizes across ancestries and TSS-distance bins (*ρ* ≤ 0.138), with direction concordance only marginally above chance. Agreement and discrimination improved for high-confidence fine-mapped variants, and Borzoi and AlphaGenome showed stronger inter-model convergence on fine-mapped variants than on nominal eQTLs, consistent with enrichment for regulatory variants whose effects are more apparent to sequence-based models. In distance-matched AUROC analyses at PIP ≥ 0.9 using PIP *<* 0.01 variants as low-PIP comparison variants, the AA high-PIP variant set yielded the highest discrimination for both Borzoi (0.837 [95% CI: 0.790–0.870]) and AlphaGenome (0.820 [0.793–0.845]). The CH-versus-NHW ordering was model-dependent: Borzoi yielded higher AUROC in NHW than CH, whereas AlphaGenome produced nearly identical CH and NHW estimates. AUROC values were lower when intermediate-PIP variants were used as comparison variants, but the AA set retained the highest discrimination. MAF-stratified sensitivity analyses attenuated some ancestry contrasts but did not eliminate the higher AA discrimination pattern. Functional annotation analysis showed that AA high-PIP variants more often overlapped chromatin accessibility and chromatin-contact annotations than NHW variants, despite lower overlap with prior eQTL and sQTL annotation catalogs.

**Conclusions:** Borzoi and AlphaGenome showed limited agreement with nominal eQTL effect sizes, but better distinguished high-confidence fine-mapped eQTLs from low-PIP variants. These results support using S2F scores as prioritization evidence for fine-mapped regulatory variants, especially promoter-proximal high-PIP variants, rather than as standalone predictors of eQTL effect size. The strongest discrimination was observed for the AA high-PIP variant set. Overall, the AA result is best interpreted as stronger separation of high-PIP variants from lower-PIP comparison variants, shaped by fine-mapping resolution, LD, the choice of comparison variants, and annotation composition.

## 1 Background

Most disease-associated variants identified by genome-wide association studies (GWAS) fall outside protein-coding sequence, and these signals are enriched in chromatin-accessible regulatory DNA [1–3]. The regulatory nature of these associations motivates approaches that predict how sequence variation alters gene regulation rather than protein structure.

Deep learning sequence-to-function (S2F) models were developed for this problem. Trained on large-scale functional genomics profiles spanning chromatin state, transcription factor binding, and gene expression, these models learn sequence-to-activity mappings that can be queried in silico to estimate cell-type-specific regulatory effects of genetic variants [4–7]. The Enformer model, developed by Avsec et al. [4], introduced transformer self-attention mechanisms to capture long-range genomic interactions up to 100 kb, substantially improving gene expression prediction accuracy. More recently, Borzoi directly models RNA-seq coverage to simultaneously predict transcription, splicing, and polyadenylation effects [6]. AlphaGenome, a recent Google DeepMind model, unifies multimodal prediction with 1 Mb sequence context and base-pair resolution, outperforming prior models on 25 of 26 variant effect prediction benchmarks [7].

Despite this methodological progress, the resources used to develop and evaluate S2F models incompletely represent global genetic diversity. The current human reference genome (GRCh38) is a biased mosaic derived primarily from a single individual, with recognized limitations in representing global genetic diversity [8]. The ENCODE and Roadmap Epigenomics consortia, which provide the majority of chromatin profiling data used in model training, profiled cell lines without systematic attention to ancestral diversity [9,10]. The GTEx consortium, a primary source of expression quantitative trait loci (eQTL) data, comprises approximately 85% European-ancestry individuals [11]. Whether S2F model evaluations remain reliable across diverse ancestry groups is therefore an open question—one made more pressing by the well-documented poor transferability of polygenic risk scores and gene expression prediction models across ancestries [12–15]. Recent work has also shown that current genomic deep learning models often fail to explain inter-individual transcriptome variation from personal genomes [16], a task whose transferability may be sensitive to ancestry-linked differences in allele frequency, haplotype background, and LD structure, even as personal-genome expression prediction remains an active modeling frontier [17].

The stakes extend beyond benchmark scores. African American, Hispanic/Latino, and other underrepresented populations face documented health disparities, and genetic tools that perform poorly for these groups risk widening rather than reducing those gaps [18,19]. Characterizing ancestry-specific performance in genomic machine learning is therefore both scientifically and ethically pressing.

Multi-ancestry cohorts provide essential resources for assessing model generalizability. The Multi-Ancestry Genomics, Epigenomics, and Transcriptomics of Alzheimer’s (MAGENTA) Project provides whole blood RNA sequencing and genome-wide genotyping from an Alzheimer disease case-control cohort across African American, Caribbean Hispanic, and Non-Hispanic White populations [20], and has supported transcriptome-wide analyses revealing both shared and population-specific genetic effects in Alzheimer disease [21]. This diversity enables direct comparison of eQTL landscapes and model behavior across groups with different allele-frequency and LD patterns.

Recent developments in fine-mapping methods attempt to address a fundamental limitation of GWAS: lead variants can tag, rather than identify, the functional variants driving an association. Statistical fine-mapping approaches such as SuSiE (Sum of Single Effects) jointly model multiple variants while accounting for LD structure, computing posterior inclusion probabilities (PIP) that quantify model support for each variant as a putative causal variant [22–24]. Previous studies have found stronger S2F model performance on fine-mapped variants than on nominal associations, consistent with fine-mapped sets being enriched for variants with direct regulatory effects [4,6,7]. Extending this evaluation across ancestries tests whether sequence-based models recognize regulatory effects in variant and haplotype backgrounds that are less represented in model-development resources, while also exposing how benchmark performance can depend on ancestry-specific LD structure and fine-mapping resolution.

Existing S2F benchmarks provide limited ancestry-stratified evidence about whether the same scoring frameworks generalize across non-European or multi-ancestry eQTL contexts. Multi-ancestry eQTL cohorts enable direct comparison within a shared tissue and analytic framework, clarifying when S2F models capture population-robust regulatory signals and when performance estimates are shaped by population genetic architecture.

Here, we address this gap through a systematic evaluation of ancestry-stratified benchmark behavior for two leading S2F models—Borzoi and AlphaGenome—using whole blood eQTL data from the MAGENTA cohort. We assess cis-eQTL prediction accuracy across African American, Caribbean Hispanic, and Non-Hispanic White populations using multiple complementary metrics: Spearman correlation and direction concordance with eQTL effect sizes, inter-model convergence, and AUROC for distinguishing SuSiE fine-mapped variants with high PIP from matched low-PIP comparison variants. Nominal eQTL analyses evaluate sampled significant SNP–gene pairs within ancestry and TSS-distance strata, while fine-mapping-based benchmarking is performed using ancestry-specific fine-mapping within each population. At stringent PIP thresholds, the AA high-PIP variant set yields higher AUROC values under this evaluation framework than the Non-Hispanic White and Caribbean Hispanic variant sets. Our findings provide a systematic assessment of S2F model behavior across diverse ancestries, clarifying where current approaches detect regulatory signal and where benchmark differences likely reflect LD structure, fine-mapping resolution, and the choice of comparison variants rather than uniformly better biological modeling in any one population.

## 2 Results

### 2.1 Dataset Overview and Fine-Mapping Summary

We analyzed whole blood eQTL summary statistics from 668 MAGENTA participants across three ancestry groups: African American (AA; *N* = 224), Caribbean Hispanic (CH; *N* = 209), and Non-Hispanic White (NHW; *N* = 235). After quality control and preprocessing, the dataset comprised 64,150 AA, 105,218 CH, and 290,846 NHW nominally significant eQTL records, involving 2,421 AA, 1,557 CH, and 3,722 NHW eGenes. After stratified sampling, model-score availability, and known TSS-distance annotation, the nominal benchmark included 93,644 scored SNP–gene records for Borzoi (AA: 29,675; CH: 27,318; NHW: 36,651) and 94,621 for AlphaGenome (AA: 30,845; CH: 29,376; NHW: 34,400). The paired nominal set available for inter-model comparison included 89,814 SNP–gene records scored by both models.

Statistical fine-mapping with SuSiE on eGenes identified high-confidence putative causal variants across all three ancestries. NHW showed the highest mean number of credible sets per eGene (0.511), followed by CH (0.357) and AA (0.272). The mean number of variants per credible set was largest in CH (53.1), compared to NHW (43.9) and AA (21.7). At PIP ≥ 0.9, NHW had the largest number of high-confidence variants (178), followed by AA (85) and CH (58). After model-score availability and TSS-distance filtering for the AUROC benchmark, PIP ≥ 0.9 positives numbered 72 AA, 51 CH, and 117 NHW pairs for Borzoi, and 79 AA, 51 CH, and 130 NHW pairs for AlphaGenome (Table S1).

### 2.2 S2F Model Scores Align More Strongly with Fine-Mapped than Nominal eQTLs

We first assessed concordance between S2F model predictions and observed eQTL effect sizes for nominally significant eQTLs, stratified by TSS distance (Figure 1a,c). Spearman correlations were consistently low across all ancestries, models, and distance bins. The strongest correlations were observed in the proximal 0–3 kb bin: Borzoi achieved *ρ* = 0.100 for AA, *ρ* = 0.052 for CH, and *ρ* = 0.085 for NHW, while AlphaGenome achieved *ρ* = 0.138, *ρ* = 0.027, and *ρ* = 0.111, respectively. Correlations declined with increasing TSS distance and were near zero beyond 35 kb, indicating that S2F model scores show detectable alignment with nominal eQTL effect sizes mainly for promoter-proximal variants. Direction concordance for nominal eQTLs was similarly modest (Figure 1c). At 0–3 kb, concordance ranged from 50.2% to 54.7% across ancestry groups and models, only marginally above the 50% chance level, and declined toward chance for distal variants. These results indicate that for nominal eQTLs—which include many LD-tagging variants—S2F model scores provide only limited predictive value for eQTL effect direction or magnitude, consistent with prior studies [16,25].

**Figure 1:**
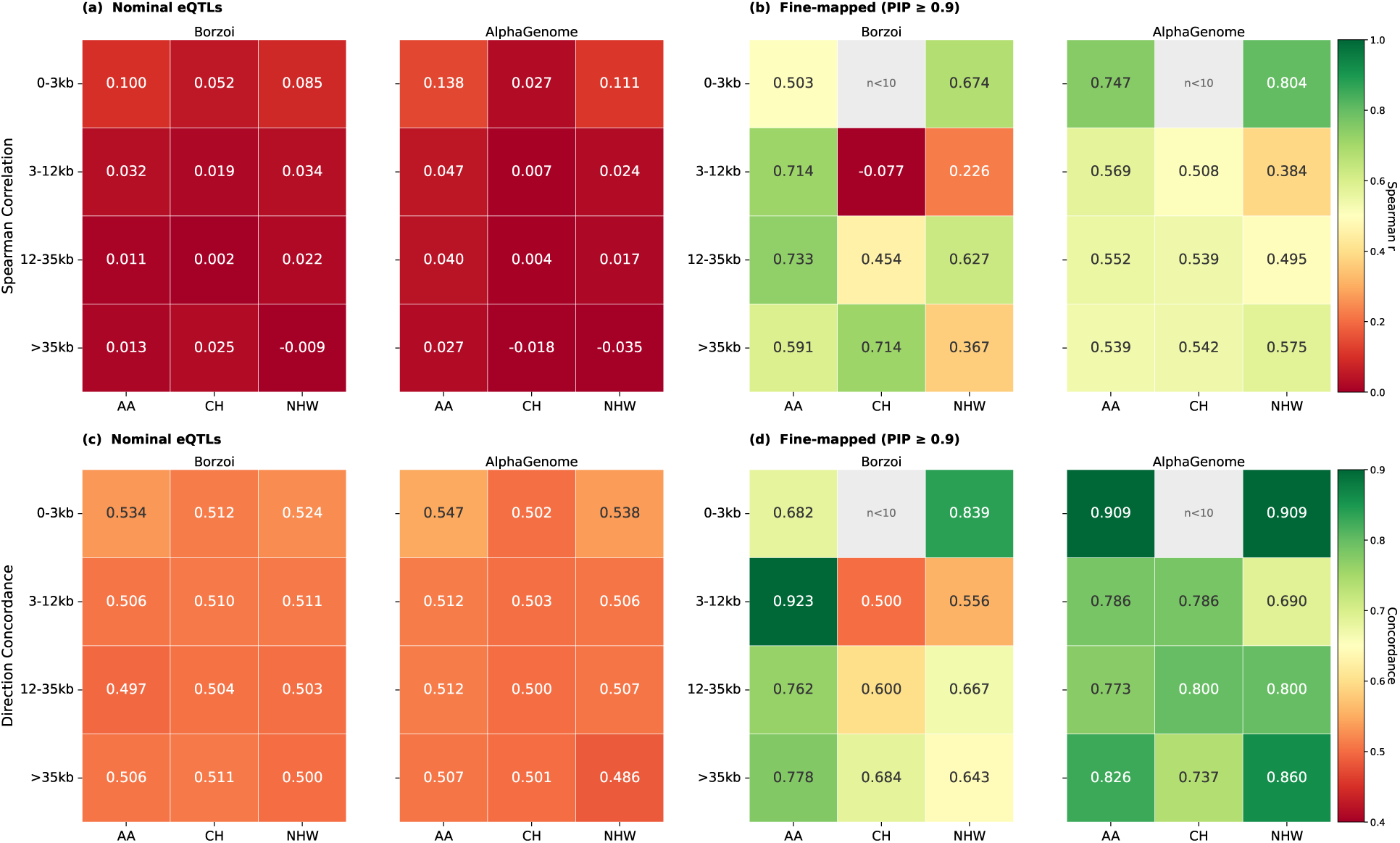
S2F model correlation and direction concordance with eQTL effect sizes, stratified by TSS distance and ancestry. **(a, b)** Spearman rank correlation between predicted variant effect scores and observed eQTL *β* coefficients for nominal eQTLs (a) and SuSiE fine-mapped variants with PIP ≥ 0.9 (b). **(c, d)** Direction concordance (proportion of variants where predicted and observed effect signs agree) for nominal eQTLs (c) and fine-mapped variants (d). All panels are stratified by distance to the transcription start site (TSS; 0–3 kb, 3–12 kb, 12–35 kb, *>*35 kb) and ancestry group (African American, Caribbean Hispanic, Non-Hispanic White), shown separately for Borzoi and AlphaGenome. Direction concordance of 0.5 indicates chance-level performance. Grey cells marked “*n <* 10” in panels (b) and (d) indicate bins excluded due to insufficient sample size (*n <* 10 fine-mapped variants); fine-mapped estimates with 10 ≤ *n <* 20 are interpreted as exploratory. The Caribbean Hispanic (CH) 0–3 kb bin contains only 4 high-confidence putative causal variants (PIP ≥ 0.9), below the threshold required for reliable correlation estimation. Counts of scored fine-mapped eQTL–gene pairs by model, ancestry, PIP threshold, and TSS-distance bin are summarized in Table S1.

Among fine-mapped variants (PIP ≥ 0.9), both correlation and concordance improved substantially (Figure 1b,d). Fine-mapped AA variants and several NHW/AlphaGenome bins showed substantially higher correlations than nominal eQTLs, with notably high values in the proximal bins (AlphaGenome: *ρ* = 0.747 for AA and *ρ* = 0.804 for NHW at 0–3 kb). Fine-mapped AA variants showed consistently high direction concordance and correlation across TSS distance bins, with higher estimates than CH and NHW in most bins. AlphaGenome showed higher direction concordance than Borzoi in all ancestry–distance combinations, except for fine-mapped AA variants in the 3–12 kb bin. CH fine-mapped variants had only 4 high-confidence putative causal variants in the 0–3 kb bin, below the minimum threshold (*n <* 10) for reliable estimation (see Figure 1 legend), and showed more variable performance in other distance bins, consistent with their smaller fine-mapped set size (*N* = 58 at PIP ≥ 0.9).

### 2.3 Borzoi and AlphaGenome Predictions Converge for Proximal Fine-Mapped Variants

To assess whether the two S2F models agree with each other on the same variants—independent of their agreement with observed eQTL effect sizes—we computed pairwise inter-model Spearman correlation and direction concordance between Borzoi and AlphaGenome predicted effect scores, stratified by TSS distance bin, ancestry, and variant class (Figure 2).

**Figure 2:**
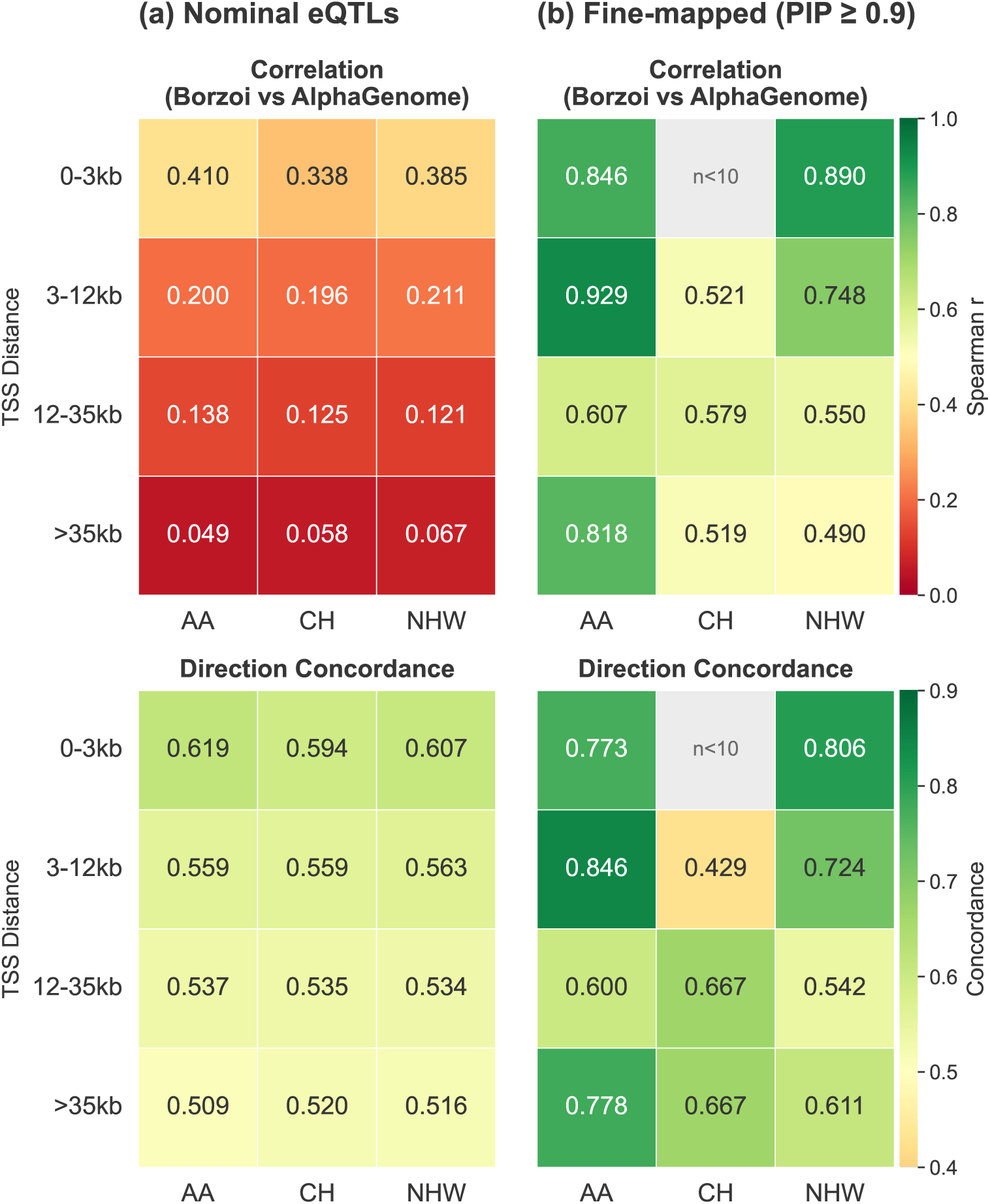
Inter-model convergence between Borzoi and AlphaGenome predictions. Heatmaps show Spearman correlation (top) and direction concordance (bottom) across TSS-distance bins and ancestry groups (AA, African American; CH, Caribbean Hispanic; NHW, Non-Hispanic White). **(a)** Nominal eQTLs. **(b)** SuSiE fine-mapped variants with PIP ≥ 0.9. Red/orange indicates lower convergence and green indicates higher convergence; concordance of 0.5 denotes chance agreement. Grey cells marked “*n <* 10” were excluded for insufficient sample size; fine-mapped estimates with 10 ≤ *n <* 20 are interpreted as exploratory. Counts of scored PIP ≥ 0.9 fine-mapped eQTL–gene pairs are summarized in Table S1; inter-model convergence uses the paired subset with scores from both Borzoi and AlphaGenome.

For nominal eQTLs, inter-model correlation was moderate for promoter-proximal variants and strongly TSS-dependent (Figure 2a, upper heatmap). Correlation ranged from 0.338–0.410 at 0–3 kb across the three ancestry groups and declined with increasing TSS distance, approaching near-zero at *>*35 kb (AA: 0.049; CH: 0.058; NHW: 0.067). Direction concordance for nominal eQTLs showed a similar gradient (Figure 2a, lower heatmap), ranging from 0.594–0.619 in the proximal bin and falling to 0.509–0.520 at *>*35 kb. These patterns indicate that the two models assign more similar scores to promoter-proximal sequence features but diverge substantially for distal variants, consistent with stronger promoter-proximal score alignment in both architectures.

For fine-mapped variants (PIP ≥ 0.9), inter-model convergence increased relative to nominal eQTLs across the available ancestry–distance strata (Figure 2b). Spearman correlations ranged from 0.49 to 0.93, with the highest point estimates observed for AA (0–3 kb: 0.846; 3–12 kb: 0.929) and NHW (0–3 kb: 0.890; 3–12 kb: 0.748). Direction concordance for fine-mapped variants was similarly elevated, with point estimates reaching 0.85 for AA in the 3–12 kb bin and 0.81 for NHW at 0–3 kb. The contrast between nominal and fine-mapped convergence is substantial: both models largely disagree on the predicted direction for nominal variants that may include LD-tagging signals, but show stronger convergence for high-confidence putative causal variants. This finding is consistent with high-PIP variants being enriched for sequence features that receive concordant S2F scores, and with the low convergence observed for nominal eQTLs reflecting the presence of LD-tagging variants rather than inherent model disagreement.

The AA high-PIP variant set showed higher inter-model convergence across several TSS distance bins and both metrics (Figure 2b), although several fine-mapped bin-level estimates were based on small subsets and should be interpreted as exploratory. At 3–12 kb, AA inter-model correlation was 0.929 (*n* = 13) compared to 0.748 for NHW and 0.521 for CH; direction concordance at the same bin was 0.846 (AA), 0.724 (NHW), and 0.429 (CH). This pattern is directionally consistent with the ancestry-stratified AUROC results and with the possibility that shorter LD blocks in African-ancestry populations can produce a more resolved fine-mapped variant set whose sequence features receive more concordant model scores, but it should be viewed as supportive rather than definitive bin-level evidence. The CH 0–3 kb bin is excluded due to insufficient sample size (*n <* 10), as noted above. AA fine-mapped variants at *>*35 kb also showed high inter-model concordance (correlation: 0.818; direction concordance: 0.778; *n* = 18). Because this distal estimate is based on a small bin, it is best treated as hypothesis-generating and could reflect fine-mapping resolution, LD and comparison-set structure, regulatory annotation composition, or model-training data composition rather than definitive evidence of AA-specific distal regulatory biology.

### 2.4 Fine-Mapped Variants Yield Higher Score-Based Discrimination than Nominal eQTLs

To assess S2F score discrimination for putative causal variants and evaluate their utility for eQTL interpretation, we performed distance-matched AUROC analysis using the absolute magnitude of the signed S2F score. The predefined high-PIP positive class (PIP ≥ *t*) was compared against two comparison sets: low-PIP variants (PIP *<* 0.01; Figure 3a) and intermediate-PIP variants (0.01 ≤ PIP *< t*; Figure 3b). AUROC was estimated over 100 bootstrap subsamples to quantify variability. Because PIP is influenced by ancestry-specific LD, allele frequency, sample size, and eQTL power, these AUROC estimates should be interpreted as discrimination under the specified choice of PIP-derived comparison variants, not as direct estimates of absolute model accuracy or experimental causality. Counts of scored positive SNP–gene pairs available for this analysis before distance matching, stratified by model, ancestry, PIP threshold, and TSS-distance bin, are summarized in Table S1.

**Figure 3:**
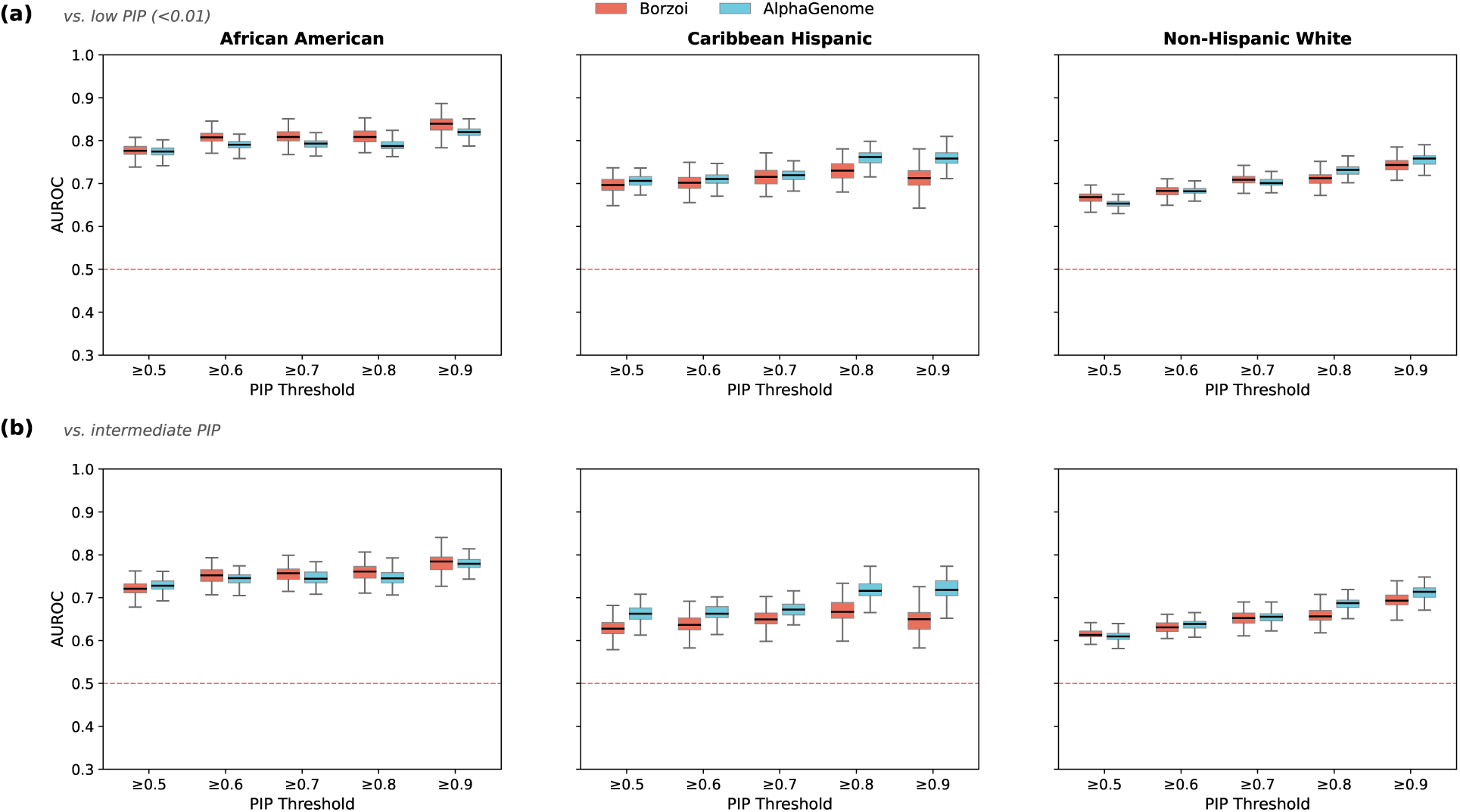
Distribution of distance-matched AUROC across 100 bootstrap iterations. **(a)** High-confidence putative causal variants (PIP ≥ *t*) vs. low-PIP comparison variants (PIP *<* 0.01). **(b)** High-confidence putative causal variants (PIP ≥ *t*) vs. intermediate-PIP variants (0.01 ≤ PIP *< t*). Panels are arranged by comparison-variant definition (rows) and ancestry (columns: AA, African American; CH, Caribbean Hispanic; NHW, Non-Hispanic White). Within each ancestry panel, Borzoi and AlphaGenome are shown as side-by-side boxplots for each SuSiE PIP threshold (0.5–0.9). Each box represents the AUROC distribution across 100 bootstrap subsamples after distance-matched sampling across TSS distance bins to control for TSS-proximity confounding. Counts of scored positive SNP–gene pairs available before distance matching are provided in Table S1; legend *n* values correspond to the matched subsets used for ROC calculation. Red dashed line indicates chance performance (AUROC = 0.5).

Against low-PIP comparison variants (PIP *<* 0.01), AUROC increased with PIP threshold for both models across all ancestry groups (Figure 3a). At PIP ≥ 0.9, the AA high-PIP variant set yielded the strongest discrimination from low-PIP comparison variants for both models, reaching 0.837 [95% CI: 0.790–0.870] for Borzoi and 0.820 [0.793–0.845] for AlphaGenome. For the non-AA groups, Borzoi separated NHW high-PIP variants from low-PIP comparison variants more strongly than CH variants (0.743 [0.708–0.773] vs. 0.714 [0.657–0.762]), whereas AlphaGenome yielded nearly identical CH and NHW estimates (0.758 [0.717–0.794] vs. 0.756 [0.729–0.783]). At lower thresholds (0.5–0.8), both models discriminated AA positives most strongly, followed by CH and then NHW, but the CH–NHW gap remained modest relative to the persistent AA-high pattern. Exploratory pairwise comparisons based on bootstrap AUROC differences supported higher AA than NHW discrimination for both Borzoi (Bonferroni-adjusted *p* = 0.0090) and AlphaGenome (*p* = 0.0060) at PIP ≥ 0.9; the AA–CH contrast was supported for Borzoi (*p* = 0.0090) but not for AlphaGenome after correction.

When the comparison set was restricted to intermediate-PIP variants (0.01 ≤ PIP *< t*; Figure 3b), AUROC values were uniformly lower, but discrimination still improved as the PIP threshold became more stringent. At PIP ≥ 0.9, Borzoi yielded AUROC = 0.779 [0.728–0.817] in AA, 0.694 [0.654–0.733] in NHW, and 0.649 [0.592–0.707] in CH; corresponding AlphaGenome values were 0.779 [0.744–0.813], 0.721 [0.670–0.767], and 0.712 [0.673–0.741]. These results support the interpretation that both models assign larger score magnitudes to high-confidence putative causal variants than to matched comparison variants, while also showing that this separation depends on the comparison set. Together, these analyses show consistently stronger discrimination for the AA fine-mapped variant set. In contrast, CH-versus-NHW ordering changed across model, PIP threshold, and comparison-set definition, so those differences should not be interpreted as a stable ancestry ranking.

### 2.5 Functional Annotation Composition of Ancestry-Specific Fine-Mapped Variants

We next asked whether ancestry-specific fine-mapped positive variants differed in their overlap with broad functional annotation categories from FILER [26]. This analysis was restricted to high-PIP positive variants and therefore tests annotation composition among fine-mapped variants rather than enrichment relative to a matched genomic background. FILER categories aggregate annotations across heterogeneous tissues and assays and were not restricted to blood-specific regulatory annotations. Sample sizes decreased with increasing PIP threshold, ranging at PIP ≥ 0.5 from 142 AA, 104 CH, and 325 NHW variants for AlphaGenome and 136 AA, 104 CH, and 309 NHW variants for Borzoi, to 76 AA, 46 CH, and 123 NHW variants for AlphaGenome and 70 AA, 46 CH, and 116 NHW variants for Borzoi at PIP ≥ 0.9 (Table S4).

Across both models, AA high-PIP variants generally showed the highest overlap with chromatin accessibility and chromatin-contact annotations, with CH usually intermediate and NHW lowest (Tables S5 and S6). At PIP ≥ 0.5, three-way chi-square tests indicated ancestry differences in accessible chromatin overlap for AlphaGenome (*q* = 0.021) and suggestive differences for Borzoi (*q* = 0.056). In both cases, AA had the highest percentage overlap (AlphaGenome: AA 70.4%, CH 63.5%, NHW 54.8%; Borzoi: AA 69.9%, CH 63.5%, NHW 55.7%). Hi-C annotations showed a similar AA-high pattern at PIP ≥ 0.5, with significant three-way chi-square tests for both AlphaGenome (*q* = 0.021) and Borzoi (*q* = 0.034): AA exceeded CH by 2.8–3.2 percentage points and NHW by 13.7–13.8 percentage points. Pairwise Fisher tests supported the larger AA–NHW differences for accessible chromatin and Hi-C at this threshold, with some Borzoi comparisons near the FDR threshold, whereas AA–CH differences were smaller and did not survive FDR correction.

The same percentage pattern persisted at PIP ≥ 0.9, but the smaller number of high-PIP variants reduced statistical support. For accessible chromatin, AA overlap remained higher than CH and NHW for AlphaGenome (AA 78.9%, CH 67.4%, NHW 65.9%; chi-square *q* = 0.393) and Borzoi (AA 78.6%, CH 67.4%, NHW 69.0%; *q* = 0.468). Hi-C showed the same descriptive ordering for AlphaGenome (AA 76.3%, CH 65.2%, NHW 59.3%; *q* = 0.186) and Borzoi (AA 77.1%, CH 65.2%, NHW 60.3%; *q* = 0.242). Thus, AA high-PIP variants consistently showed higher overlap percentages for these annotations, but the PIP ≥ 0.9 comparisons should be interpreted descriptively rather than as statistically significant after FDR correction. Additional chromatin-contact annotations showed concordant but generally weaker patterns: overlap with IM-PET predicted long-range chromatin interactions was higher in AA than in CH and NHW at PIP ≥ 0.5 for both AlphaGenome (48.6%, 43.3%, and 34.5%) and Borzoi (49.3%, 43.3%, and 35.6%), while overlap with pcHi-C promoter-centered chromatin contacts was high in all three ancestries and showed smaller differences. In contrast, prior QTL catalog overlap followed the opposite direction. In our FILER summary, a high-PIP variant was counted as overlapping the eQTL category if it overlapped at least one previously reported significant eQTL association represented in FILER. Because major QTL resources contributing to such catalogs, including GTEx v8 and the eQTL Catalogue, are dominated by European-ancestry samples, ancestry differences in overlap may partly reflect the discovery history and sample composition of prior QTL studies [11,27,28]. At PIP ≥ 0.5, three-way chi-square tests indicated strong ancestry differences in eQTL annotation overlap for both AlphaGenome (AA: 83.1%; CH: 96.2%; NHW: 95.1%; *q* = 1.86 × 10^−4^) and Borzoi (AA: 84.6%; CH: 96.2%; NHW: 96.8%; *q* = 5.75 × 10^−5^). Here, both CH and NHW had higher overlap than AA, with AA lower by 11.6–13.1 percentage points relative to CH and by 12.0–12.2 percentage points relative to NHW. This pattern persisted descriptively at PIP ≥ 0.9 (AlphaGenome: AA 81.6%, CH 97.8%, NHW 93.5%; Borzoi: AA 84.3%, CH 97.8%, NHW 95.7%). sQTL annotations showed a similar contrast at lower thresholds, with AA overlap lower than CH and NHW at PIP ≥ 0.5 for both models. Together, these results suggest that AA high-PIP variants frequently overlapped regulatory annotations, particularly chromatin accessibility and chromatin-contact annotations, while being less represented in prior, European-ancestry-dominated QTL annotation catalogs.

## 3 Discussion

In this study, we evaluated two long-context sequence-to-function deep learning models, Borzoi and AlphaGenome, for regulatory variant-effect prediction using whole blood eQTLs across three ancestrally diverse populations. Our key findings are: (1) both models show weak agreement with nominal eQTL effect sizes but meaningful discriminative power for fine-mapped putative causal variants; (2) the AA high-PIP variant set yields the highest benchmark discrimination under our evaluation framework; and (3) ancestry-stratified benchmark patterns likely reflect a combination of LD architecture, fine-mapping resolution, functional-annotation context, and comparison-set definition, rather than a single ancestry-intrinsic property of the models.

### Fine-mapped variants show stronger S2F score alignment

Spearman correlations between model predictions and observed eQTL effect sizes were consistently low for nominally significant eQTLs (*ρ* ≤ 0.138 across ancestry and distance strata), consistent with prior benchmarking showing that S2F models have limited power to rank eQTL effect sizes in population-level studies [7,16,25,29]. This low correlation should not necessarily be interpreted as a failure of the models to identify functional variants. S2F models estimate sequence-derived regulatory impact learned from functional genomics data, whereas nominal eQTL effect sizes reflect statistical association signals that can be distributed across many correlated variants through LD, and context-specific regulatory effects. Thus, nominal eQTL sets likely contain a mixture of functionally impactful variants and LD-tagging variants with little direct sequence-level effect, attenuating variant-level correlations between model scores and observed association estimates. Both models were also trained on a linear reference genome and do not explicitly represent population-level haplotype diversity or LD structure, which may further decouple sequence-function predictions from association-based eQTL estimates [25]. When restricted to fine-mapped putative causal variants (PIP ≥ 0.9), however, both Spearman correlation and AUROC rose substantially. This contrast suggests that fine-mapping enriches the benchmark for variants that receive larger or more concordant S2F scores, while nominal eQTL benchmarks dilute this signal with LD-linked variants that are statistically associated but not necessarily causal.

This interpretation is further supported by the inter-model convergence analysis (Figure 2). Borzoi and AlphaGenome are not fully independent in design: both are long-context sequence-to-function models that use convolutional and transformer components. However, they differ in their implementation, training objectives, and output spaces. Borzoi extends the Enformer modeling framework to predict RNA-seq coverage and related functional genomic tracks across a 524 kb sequence window, whereas AlphaGenome uses a 1 Mb U-Net-inspired multimodal architecture to predict base-pair-resolution regulatory tracks, splice-related outputs, and contact maps [6,7]. Against this background, the convergence pattern is informative: the two models largely disagree on predicted effect directions for nominal eQTL variants (direction concordance near chance at *>*35 kb), but show stronger convergence for fine-mapped putative causal variants (direction concordance up to 0.85 for AA, 0.81 for NHW at proximal bins). Because some fine-mapped bin-level estimates are based on small subsets, these maxima should be interpreted as descriptive point estimates rather than standalone evidence. This between-model convergence suggests that the score pattern at fine-mapped loci is less likely to reflect a peculiarity of one model or scoring strategy alone, particularly when the benchmark is enriched for promoter-proximal putative causal variants.

The strong TSS-distance dependence we observe—with inter-model correlation falling from ∼0.34–0.41 at 0–3 kb to ∼0.05–0.07 at *>*35 kb for nominal eQTLs—is consistent with stronger promoter-proximal score alignment by both models. Although Borzoi’s 524 kb input window and AlphaGenome’s 1 Mb window are architecturally capable of capturing long-range regulatory elements, both models predict promoter-proximal regulatory activity better. Our findings therefore suggest that S2F scores are more informative for genes with proximal regulatory architecture, and that caution is warranted when applying these scores to distal enhancer-driven or cell-type-specific regulatory contexts [29]. At the same time, this distance-dependent decline was much weaker among fine-mapped variants. Fine-mapping likely enriches the evaluated set for variants that are more directly involved in regulation, rather than variants that simply tag a causal signal through LD. Thus, even for variants located farther from the TSS, the two models may still converge on their predicted effects because the fine-mapped set contains a higher fraction of putative causal regulatory perturbations.

### Ancestry-dependent benchmark behavior

The AA fine-mapped variant set showed stronger score alignment across several complementary analyses, not only in AUROC. In the correlation and direction-concordance analyses, AA fine-mapped variants showed consistently high estimates across TSS-distance bins, with higher values than CH and NHW in most bins. The inter-model convergence analysis provided a second line of evidence: Borzoi and AlphaGenome showed high agreement for AA across several TSS-distance bins, including at 3–12 kb, where inter-model Spearman correlation reached 0.929 and direction concordance reached 0.846. Because several fine-mapped bin-level estimates are based on small subsets, these heatmap results should be interpreted as supportive and exploratory at the bin level rather than as definitive evidence from any single heatmap bin. Together, these results support a broader pattern of higher score alignment for the AA fine-mapped set than for the corresponding CH or NHW fine-mapped sets under this benchmark framework.

AUROC was used to assess how well each model’s score magnitude separated high-PIP putative causal variants from low-PIP or intermediate-PIP comparison variants within each ancestry group. The clearest ancestry contrast was the AA-high pattern, which appeared across both models and comparison-set definitions. By contrast, CH-versus-NHW ordering changed across model, PIP threshold, and comparison-set definition, so those differences should not be interpreted as a stable ancestry ranking.

The AA-high discrimination pattern is unlikely to be explained by a single factor, but the most direct explanation is fine-mapping resolution. Although the same PIP threshold was applied across ancestries, AA credible sets were smaller on average (21.7 variants per credible set) than NHW (43.9) or CH (53.1), suggesting that posterior mass was concentrated among fewer candidate variants. A more resolved high-PIP positive class would be easier for sequence-derived scores to separate from low- or intermediate-PIP comparison variants. In longer-LD or admixed populations, comparison variants may remain more tightly correlated with the inferred fine-mapped signal, making them harder for sequence-based scores to rank below high-PIP variants [30]. The attenuation of AA–non-AA AUROC gaps when intermediate-PIP variants were used as comparison variants further indicates that comparison-set composition contributes to this separation, and MAF-stratified sensitivity analyses showed that allele-frequency composition alone does not eliminate the stronger AA discrimination (Figure S1 and Table S3). Thus, the observed ancestry pattern should be interpreted primarily as a property of this fine-mapping-based benchmark rather than as evidence of uniformly better biological modeling in AA.

### Functional annotation overlap provides regulatory context

The FILER annotation analysis provides complementary context for interpreting the ancestry-stratified benchmark. AA fine-mapped variants were not annotation-poor overall: across both S2F models, they more often overlapped chromatin accessibility and chromatin-contact annotations than NHW variants, with concordant patterns for Hi-C, IM-PET, and pcHi-C categories. Because this was a positive-variant composition analysis rather than a matched enrichment test, these overlaps should be treated as contextual support rather than a causal explanation for the AUROC pattern. The FILER chromatin-contact annotations contributing to these patterns were heterogeneous, including lymphoblastoid and hematopoietic cell lines such as GM12878 and K562 as well as primary immune-cell datasets. In an audit of overlapping Hi-C, IM-PET, and pcHi-C annotation metadata, we did not identify HeLa-derived contact tracks; therefore, the AA-high chromatin-contact pattern should be interpreted as overlap with heterogeneous 3D regulatory maps rather than evidence for a specific HeLa- or cell-line-driven mechanism. At the same time, AA high-PIP variants showed lower overlap with prior eQTL and sQTL annotation catalogs than CH and NHW variants. We interpret this contrast cautiously: prior QTL annotations reflect the ancestry composition, tissue coverage, sample size, and LD structure of existing resources, and therefore should not be treated as a direct measure of intrinsic regulatory function. Nevertheless, the opposing directions of chromatin annotation overlap and QTL catalog overlap are consistent with the broader concern that current regulatory annotation resources are not ancestry-neutral [27,28]. These results suggest that ancestry differences in S2F benchmarks may reflect both fine-mapping resolution and the uneven representation of regulatory evidence across annotation modalities.

### Practical interpretation

These results support using Borzoi and AlphaGenome scores as one layer of evidence for prioritizing fine-mapped regulatory variants, not as standalone predictors of eQTL effect size. Confidence is strongest for promoter-proximal and high-PIP variants, where model agreement and AUROC improved relative to nominal eQTLs. For distal variants, nominal eQTLs, or ancestry comparisons, S2F scores should be interpreted together with fine-mapping uncertainty, LD structure, the choice of comparison variants, and functional annotation context. In practice, ancestry-stratified S2F benchmarks should therefore use population-specific fine-mapping and matched comparison sets rather than assuming that a score threshold calibrated in one ancestry or variant class transfers directly to another.

### Limitations

Several limitations should be noted. First, MAGENTA is modest in size (*N* = 224–235 per ancestry), limiting eQTL power and fine-mapping resolution, particularly for CH high-PIP variants. Fine-mapping results depend on LD reference quality, association power, prior assumptions, and PIP calibration; consequently, PIP thresholds should be interpreted as statistical support rather than experimental proof of causality. Second, whole blood is a heterogeneous tissue, and residual variation in cell-type composition may obscure cell-type-specific regulatory effects even after covariate adjustment. Because participants were ascertained through an Alzheimer disease case-control study, residual disease-related expression effects may also influence blood eQTL estimates despite inclusion of AD status as a covariate.

Third, although the population labels used in this study are correlated with genetic ancestry in this cohort, they are self-identified population categories analyzed with genetic-ancestry covariates and should not be interpreted as discrete biological boundaries. Our AUROC analyses used absolute score magnitude and PIP-derived comparison variants; exploratory pairwise ancestry comparisons were post hoc tests derived from the bootstrap AUROC distributions rather than prespecified inference models. These tests quantify variability from the distance-matched resampling procedure and do not capture uncertainty in eQTL discovery, LD estimation, or SuSiE fine-mapping. The sampled nominal benchmark remains a noisy association-based reference and should not be interpreted as a direct catalogue of causal regulatory variants. Fourth, the FILER overlap analysis is descriptive rather than a matched enrichment test, and regulatory annotation catalogs may overlap conceptually or empirically with resources used during model development. Finally, these results are limited to blood eQTLs and require validation in additional tissues, larger multi-ancestry cohorts, cell-type-specific regulatory maps, and experimental perturbation assays.

## 4 Conclusions

This study provides a multi-ancestry benchmark of Borzoi and AlphaGenome for whole blood eQTL interpretation across African American, Caribbean Hispanic, and Non-Hispanic White populations. Both models show weak agreement with nominal eQTL effect sizes but meaningful discriminative power for high-confidence SuSiE fine-mapped putative causal variants. At PIP ≥ 0.9 against distance-matched PIP *<* 0.01 comparison variants, the AA high-PIP variant set yields the highest AUROC for both models (Borzoi: 0.837; AlphaGenome: 0.820), whereas the CH-versus-NHW ordering is model-dependent. Chromatin accessibility and chromatin-contact overlap provide supportive context for this pattern, while the comparison-set analyses indicate that LD structure, fine-mapping resolution, and the choice of comparison variants also shape benchmark separability—the separation of high-PIP variants from lower-PIP comparison variants.

These findings support using S2F scores as prioritization evidence for fine-mapped regulatory variants, especially promoter-proximal high-PIP variants, rather than as standalone predictors of eQTL effect size. Model utility depends on variant proximity to the TSS, population LD architecture, comparison-set definition, and annotation composition. The higher AUROC observed for the AA high-PIP variant set should therefore be interpreted primarily as stronger separation under this benchmark, rather than as evidence that the models uniformly represent African American regulatory biology better than other populations. Future work should test whether S2F models trained, fine-tuned, or calibrated with more ancestrally diverse genomic and functional data can clarify and potentially reduce ancestry-dependent benchmark behavior.

## 5 Methods

### 5.1 Study Cohort

Whole blood eQTL summary statistics were obtained from the Multi-Ancestry Genomics, Epigenomics, and Transcriptomics of Alzheimer’s (MAGENTA) cohort [20], a multi-site study ascertained at the John P. Hussman Institute for Human Genomics at the University of Miami Miller School of Medicine, North Carolina A&T State University, and Case Western Reserve University. The cohort comprises participants of three self-identified ancestries: African American (AA; *N* = 224), Caribbean Hispanic (CH; *N* = 209), and Non-Hispanic White (NHW; *N* = 235). Participants were enrolled as part of the Alzheimer’s Disease Sequencing Project (ADSP) Follow-up Study, including clinically adjudicated Alzheimer’s disease (AD) cases (onset *>* 65 years) and cognitively normal controls (age at examination *>* 65 years).

### 5.2 Genotyping and Quality Control

All participants were genotyped using the Illumina MEGA v2 or v3 multi-ancestry array and imputed against TOPMed reference panels (build GRCh38), with ancestry-matched panels applied per population group [31]. Variants were retained if directly genotyped or imputed with quality score *R*^2^ *>* 0.8, and subsequently filtered to exclude variants with minor allele count (MAC) *<* 10, Hardy–Weinberg equilibrium *P <* 10^−8^, or missingness *>* 0.1; sample-level missingness *>* 0.1 was similarly applied. Pairwise relatedness was estimated using KING [32], and one sample from each pair with kinship coefficient ≥ 0.0625 was removed, prioritizing retention of cognitively normal elderly individuals, AD cases, and carriers of rare APOE genotypes. Population substructure was assessed by principal components analysis (PCA) using the flashpca package [33] on LD-pruned variants generated with PLINK2.0 (*r*^2^ *<* 0.1, window 50, step 10) [34], and PCA outliers were excluded. All genotyping quality control was implemented following the ADSP xQTL protocol (v0.1.1).

### 5.3 RNA Processing and Expression Quantification

RNA extraction, library preparation, and sequencing of whole blood samples have been previously described [20]. Briefly, RNA was extracted from previously frozen whole blood, quality controlled to a minimum RNA integrity number (RIN) of 5, and prepared using the NuGEN Universal Plus mRNA-Seq protocol with globin and ribosomal depletion. Sequencing was performed at 125 bp paired-end on Illumina HiSeq3000. Reads were aligned to GRCh38 using STAR (v2.7.10a) [35], quality controlled using Picard, and quantified using RSEM (v1.3.3) [36]. Low-expression genes—defined as TPM in the bottom 10th percentile in more than 20% of samples, or read counts below 6 in fewer than 20% of samples—were excluded. Sample-level expression outliers were identified and removed based on relative log expression (RLE), hierarchical clustering, and D-statistics. Retained expression values were normalized using the Trimmed Mean of M-values (TMM) method [37].

### 5.4 Covariate Construction and eQTL Mapping

The full covariate set for cis-eQTL mapping comprised age at examination, sex, AD case/control status, 1–3 principal components of genetic ancestry, CIBERSORTx-derived cellular heterogeneity estimates [38], and hidden confounding factors derived from Marchenko–Pastur PCA as implemented in the ADSP xQTL protocol.

Cis-eQTL mapping was performed using TensorQTL [39], with cis-windows defined by topologically associated domain boundaries (TADB) for each gene as implemented in the ADSP FunGen-xQTL protocol (v0.1.1; https://github.com/StatFunGen/xqtl-protocol). AD case/control status was included as a covariate in all nominal eQTL analyses. Significance was assessed using a per-gene *q*-value threshold of *<* 0.05 combined with an individual variant nominal *p*-value *<* 2.02 × 10^−5^, both thresholds adopted from the eQTLGen Consortium [40]. All mapping was performed independently within each ancestry group. The resulting dataset comprised 290,846 NHW, 105,218 CH, and 64,150 AA cis-eQTL records. For nominal S2F benchmarking, significant SNP–gene pairs were stratified by ancestry group and TSS-distance bin. To prevent very large strata from dominating the benchmark, we sampled up to 10,000 records per ancestry group and TSS-distance bin before scoring and downstream analysis; smaller strata were retained in full.

### 5.5 eQTL Preprocessing and Annotation

eQTL summary statistics were in GRCh38/hg38 coordinates and annotated with the distance to the transcription start site (TSS) of each target gene, derived from Ensembl gene annotations. Variants were classified into four TSS distance bins: 0–3 kb (proximal; 4.0% of eQTLs), 3–12 kb (9.4%), 12–35 kb (17.7%), and *>*35 kb (distal; 68.0%). Scored variant records were represented on the GRCh38/hg38 forward strand, with the reference allele defined by the hg38 reference genome and the eQTL beta measuring the alternate-allele effect. Direction-based analyses therefore compared signed model predictions for the explicit reference-to-alternate substitution with the corresponding alternate-allele eQTL beta.

### 5.6 SuSiE Fine-Mapping

Statistical fine-mapping was performed on eGenes (genes with at least one significant eQTL) using the Sum of Single Effects (SuSiE) model [22,23], independently for AA, CH, and NHW populations. For each eGene, we included all variants in the cis-regulatory region, which was defined in Section 5.4, retaining variants with non-missing summary-statistic *z* scores computed as *β/*SE. SuSiE RSS fine-mapping used the MAGENTA ancestry-specific LD matrix for variants in the corresponding cis-regulatory region and was run with *L* = 10 single-effect components. Credible sets used 95% coverage and SuSiE’s default purity filter (minimum absolute correlation of 0.5; equivalently, reported `cs_min_r2` ≥ 0.25). The aggregate fine-mapping summaries analyzed here were generated from the original run without residual-variance estimation; prior variance was estimated by SuSiE, and all other SuSiE parameters were left at defaults. SuSiE computes a posterior inclusion probability (PIP) for each variant, quantifying model support for that variant as a putative causal variant given the observed association data and LD structure.

### 5.7 Sequence-to-Function Models

We evaluated two recent long-context deep learning sequence-to-function (S2F) models. **Borzoi** [6] is a convolutional–transformer model that predicts strand-specific RNA-seq coverage across a 524 kb genomic window, trained on multi-species RNA-seq, CAGE, and chromatin accessibility data. Analyses used the Borzoi human Replicate 3 model (borzoi v1.0.1). **AlphaGenome** [7] accepts 1 Mb of genomic sequence and jointly predicts RNA-seq, CAGE, chromatin accessibility, and other regulatory signals at base-pair resolution; analyses used alphagenome v0.5.0. For both models, gene-expression variant effects were quantified using the same RNA-seq scoring scheme recommended for AlphaGenome gene-expression variant scoring. Predictions were generated separately for the reference and alternate allele sequences, restricted to the whole-blood RNA-seq output for the target gene, and summarized across the annotated exons of that gene. The signed score was then computed as the alternate-minus-reference log-fold change in predicted gene expression:

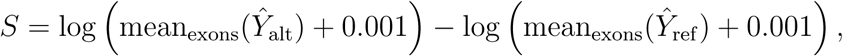

Where *Ŷ*_alt_ and *Ŷ*_ref_ denote predicted RNA-seq coverage for the alternate and reference sequences, respectively. Positive values indicate higher predicted expression for the alternate allele. In the scored output tables, this signed score corresponds to Borzoi `logMeanSED` and the AlphaGenome whole-blood RNA-seq effect score stored as `raw_score`; both columns were generated using the same exon-masked RNA-seq expression scoring definition above. Both models use the GRCh38 human reference genome as input. Due to their differing context window sizes, variants located beyond 262 kb (Borzoi) or 500 kb (AlphaGenome) from the target gene TSS could not be scored and were excluded from downstream analyses.

### 5.8 Spearman Correlation and Direction Concordance

To assess how well S2F model scores recapitulate the observed direction and magnitude of eQTL effects, we computed two complementary metrics—Spearman’s rank correlation and sign concordance (direction concordance)—applied separately to two variant sets: (1) nominally significant eQTLs and (2) SuSiE fine-mapped variants (PIP ≥ 0.9).

For **nominal eQTLs**, Spearman’s rank correlation was computed between the signed predicted variant effect score and the observed eQTL effect size (*β* coefficient) within the sampled nominal benchmark for each ancestry group. Nominal eQTLs include a mixture of putative causal and LD-tagging variants, providing a realistic but noisy benchmark. For **fine-mapped variants**, the same correlation analysis was applied to high-confidence putative causal variants (PIP ≥ 0.9), which are enriched for regulatory signal and therefore provide a higher signal-to-noise benchmark. In both cases, analyses used the signed alternate-minus-reference, exon-masked whole-blood RNA-seq S2F effect score described above (Borzoi `logMeanSED`; AlphaGenome whole-blood RNA-seq score stored as `raw_score`) and were stratified by TSS distance bin (0–3 kb, 3–12 kb, 12–35 kb, *>*35 kb) and ancestry group to characterize performance heterogeneity.

Direction concordance—the proportion of variants for which the sign of the predicted effect score matches the sign of the observed *β*—was computed for both variant sets across all ancestry groups and TSS distance bins. A concordance of 50% indicates chance-level performance.

Statistically significant differences in Spearman correlation between ancestry pairs were tested using Fisher’s *r*-to-*z* transformation [41].

### 5.9 Distance-Matched AUROC Analysis

To evaluate whether S2F model score magnitudes can discriminate high-PIP putative causal variants from lower-PIP comparison variants, we performed a receiver operating characteristic (ROC) analysis using ancestry-specific SuSiE fine-mapping. This approach was adapted from the variant effect prediction benchmarking framework described by Avsec et al. [7], but the positive class was fixed to the predefined high-PIP benchmark rather than redefined from newly discovered positives in the expanded prediction run. Specifically, positives were taken from the original scored SuSiE table and required ancestry assignment (AA, CH, or NHW), nonzero credible-set membership, a known TSS-distance bin, and PIP ≥ *t*, where *t* ∈ {0.5, 0.6, 0.7, 0.8, 0.9}. At PIP ≥ 0.5, this yielded 596 unique positive variant–gene pairs (AA: 147, CH: 113, NHW: 336), all of which were recovered in the corresponding per-gene SuSiE files used for resampling.

The background set was rebuilt from the per-gene all-variant SuSiE outputs. Low-PIP comparison variants were defined as variants with PIP *<* 0.01 and were sampled within strata defined by ancestry, TSS-distance bin, and minor allele frequency (MAF) bin, targeting 100 low-PIP comparison variant–gene pairs per positive stratum. At PIP ≥ 0.5, this produced 59,600 unique low-PIP comparison variant–gene pairs overall (AA: 14,700; CH: 11,300; NHW: 33,600). We additionally retained intermediate-PIP comparison variants (0.01 ≤ PIP *< t*) for an alternative comparison-set analysis.

Because TSS distance strongly confounds S2F model performance—proximal variants are systematically better predicted than distal ones—we controlled for this confounder using a distance-matching approach over variant–gene pairs. The log_10_(distance to TSS) was divided into 10 equal-width bins. Within each bin and bootstrap iteration, positive pairs were retained and comparison pairs were sampled with replacement to match the number of positives in that bin, yielding a distance-balanced dataset. AUROC was computed using the absolute magnitude of the same signed exon-masked RNA-seq S2F effect score—Borzoi `logMeanSED` and the corresponding AlphaGenome whole-blood RNA-seq score stored as `raw_score`—rather than signed effect direction. This procedure was repeated 100 times with different random seeds, and the mean AUROC with 95% confidence interval was reported. As an exploratory post hoc comparison, pairwise ancestry differences in AUROC were evaluated from the stored bootstrap AUROC distributions using empirical two-sided tests on bootstrap differences, followed by Bonferroni correction within each model and comparison-set definition. These tests assess variation from the distance-matched resampling procedure and do not account for uncertainty in eQTL discovery, LD estimation, or SuSiE fine-mapping. For sensitivity analysis, we repeated the same distance-matched bootstrap within three MAF strata (MAF *<* 0.05, 0.05 ≤ MAF *<* 0.2, and MAF ≥ 0.2).

### 5.10 Functional Annotation Overlap Analysis

To contextualize the regulatory features of high-PIP fine-mapped variants, we annotated variants with a FILER2 blood-oriented subset in which all tracks were labeled with Tissue category = Blood (FILER-BED Schema V2, 2024-11-06; hg38/GRCh38, including tracks lifted to hg38) using FILER_giggle v0.6.3fsbv [26]. Because scored high-PIP variant sets differed slightly between Borzoi and AlphaGenome after model-score availability filtering, annotation analyses were performed separately for the variants available to each model. Variants were evaluated at cumulative PIP thresholds of ≥ 0.5, ≥ 0.6, ≥ 0.7, ≥ 0.8, and ≥ 0.9.

For downstream summaries, overlapping annotations were collapsed into broad FILER functional categories, such as accessible chromatin, histone ChIP-seq peaks, TF ChIP-seq peaks, chromatin interactions, and QTLs. Thus, category-level annotations reflect heterogeneous blood and immune-cell assays, including both primary cells and cell lines, rather than a single whole-blood or cell-type-specific regulatory map. For chromatin-contact categories emphasized in the Results, we additionally inspected source-track metadata for overlapping Hi-C, IM-PET, and pcHi-C annotations to summarize the contributing cell lines, cell types, and source resources. For each category, overlap counts were collapsed to binary indicators, such that a variant was considered annotated if it overlapped at least one FILER record in that category. For QTL categories, this binary indicator asked whether a high-PIP variant overlapped at least one previously reported significant eQTL or sQTL association represented in FILER. For each model, PIP threshold, and annotation category, we compared annotation overlap fractions across AA, CH, and NHW variants using 3×2 chi-square tests. Pairwise ancestry comparisons were performed using two-sided Fisher’s exact tests. Benjamini–Hochberg FDR correction was applied separately within each model and PIP threshold. This analysis compares annotation overlap among high-PIP variants only and should not be interpreted as formal enrichment relative to all tested variants or a matched genomic background.

## Acknowledgements

This work was supported by grants from the National Institute on Aging R01AG070935 (Bush, Griswold), U19AG074865 (Pericak-Vance, Byrd, Bush, Haines, Kunkle, Reitz, Vance), and U54AG052427 (Wang, Schellenberg). This work made use of the High Performance Computing Resource in the Core Facility for Advanced Research Computing at Case Western Reserve University.

## Data Availability

Derived summary resources required to reproduce the manuscript figures and tables will be made publicly available before publication.

## Code Availability

Analysis code used for model scoring, summary analyses, and manuscript figure and table generation is available at https://github.com/Xinyu-Sun/s2f_snp_eqtl_analysis_code.

