## Supplemental Material for "Evaluating sequence-to-function deep learning models for ancestry-stratified regulatory variant effect prediction using multi-ancestry blood eQTLs"

### S1 Supplementary Figures

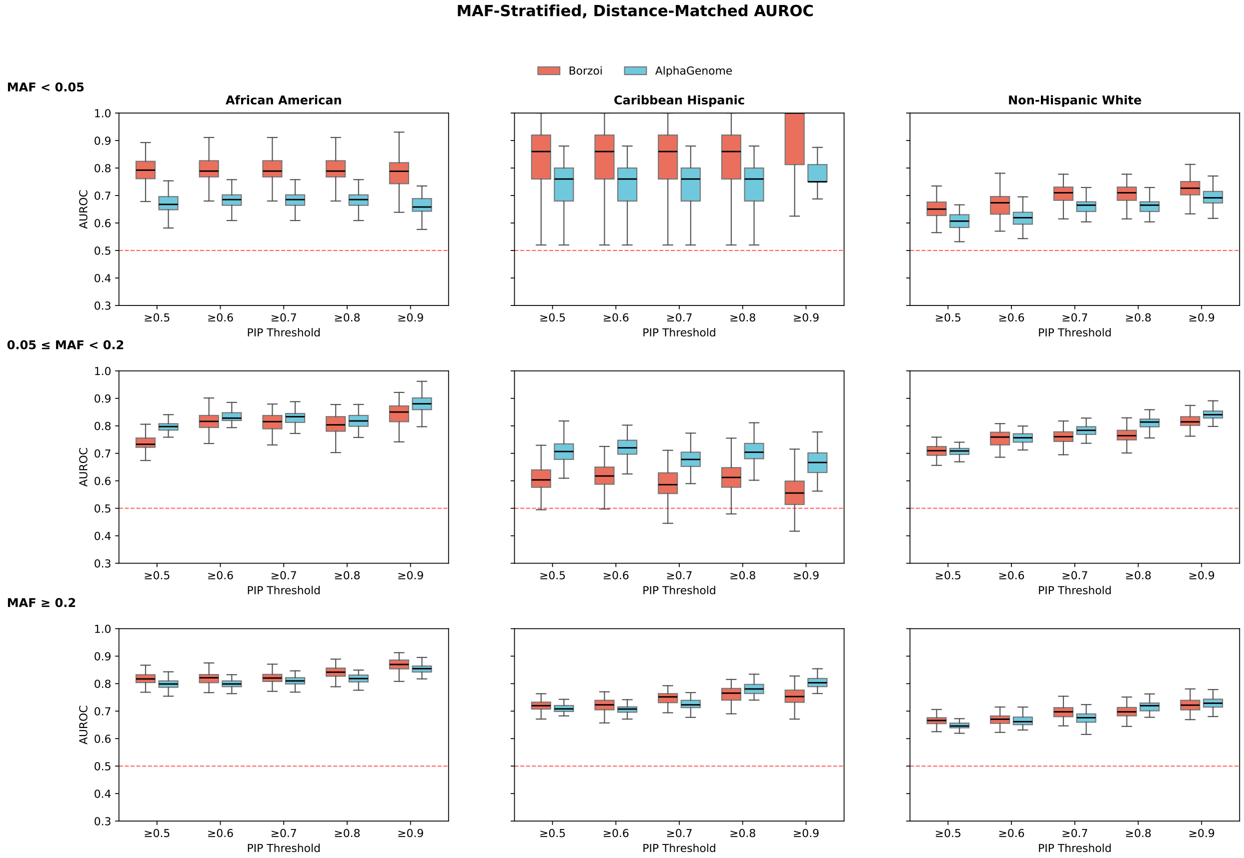


Figure S1: **MAF-stratified, distance-matched AUROC across ancestry groups and models.** AUROC was computed separately within each MAF stratum for African American (AA), Caribbean Hispanic (CH), and Non-Hispanic White (NHW) variants, using the same distance-matched framework as in the primary AUROC analysis. Columns denote ancestry groups and rows denote MAF strata. Within each panel, Borzoi and AlphaGenome are shown as side-by-side boxplots for each SuSiE PIP threshold (0.5–0.9), and each box summarizes 100 bootstrap iterations. The red dashed line indicates chance performance (AUROC $= 0.5$). Estimates in the MAF $< 0.05$ stratum, particularly for CH, should be interpreted cautiously because the number of positive variants is small.

### S2 Supplementary Tables

Table S1: **Supplementary Table S1. Counts of scored fine-mapped eQTL–gene pairs by model, ancestry, PIP threshold, and TSS-distance bin for manuscript benchmark analyses.** Counts are shown for the deduplicated manuscript positive benchmark after restricting to positive pairs with a valid recomputed TSS distance and a model score. Rows with PIP $\geq$ 0.9 correspond to the high-confidence fine-mapped benchmark used in the fine-mapped panels of Figure 1 and the PIP $\geq$ 0.9 AUROC analyses in Figure 3. Inter-model convergence in Figure 2 uses the paired subset with scores from both Borzoi and AlphaGenome, so cell counts may be equal to or smaller than the per-model counts shown here. The $n$ values shown in AUROC figure legends are smaller than these totals because positive and comparison pairs were subsequently distance-matched within log-distance bins before ROC calculation. Fine-mapped heatmap estimates based on $10 \leq n < 20$ are interpreted as exploratory; cells with $n < 10$ were omitted.

| **Model** | **Group** | **PIP** | **Total** | **0–3 kb** | **3–12 kb** | **12–35 kb** | **>35 kb** |
|---|---|---|---|---|---|---|---|
| Borzoi | AA | $\geq$ 0.5 | 140 | 41 (29.3%) | 25 (17.9%) | 40 (28.6%) | 34 (24.3%) |
| Borzoi | AA | $\geq$ 0.6 | 124 | 35 (28.2%) | 24 (19.4%) | 36 (29.0%) | 29 (23.4%) |
| Borzoi | AA | $\geq$ 0.7 | 106 | 32 (30.2%) | 21 (19.8%) | 28 (26.4%) | 25 (23.6%) |
| Borzoi | AA | $\geq$ 0.8 | 93 | 28 (30.1%) | 18 (19.4%) | 24 (25.8%) | 23 (24.7%) |
| Borzoi | AA | $\geq$ 0.9 | 72 | 22 (30.6%) | 13 (18.1%) | 19 (26.4%) | 18 (25.0%) |
| Borzoi | CH | $\geq$ 0.5 | 113 | 15 (13.3%) | 28 (24.8%) | 28 (24.8%) | 42 (37.2%) |
| Borzoi | CH | $\geq$ 0.6 | 100 | 14 (14.0%) | 23 (23.0%) | 24 (24.0%) | 39 (39.0%) |
| Borzoi | CH | $\geq$ 0.7 | 82 | 8 (9.8%) | 20 (24.4%) | 24 (29.3%) | 30 (36.6%) |
| Borzoi | CH | $\geq$ 0.8 | 66 | 6 (9.1%) | 18 (27.3%) | 20 (30.3%) | 22 (33.3%) |
| Borzoi | CH | $\geq$ 0.9 | 51 | 4 (7.8%) | 14 (27.5%) | 15 (29.4%) | 18 (35.3%) |
| Borzoi | NHW | $\geq$ 0.5 | 313 | 75 (24.0%) | 80 (25.6%) | 68 (21.7%) | 90 (28.8%) |
| Borzoi | NHW | $\geq$ 0.6 | 236 | 56 (23.7%) | 58 (24.6%) | 51 (21.6%) | 71 (30.1%) |
| Borzoi | NHW | $\geq$ 0.7 | 184 | 48 (26.1%) | 44 (23.9%) | 38 (20.7%) | 54 (29.3%) |
| Borzoi | NHW | $\geq$ 0.8 | 149 | 41 (27.5%) | 36 (24.2%) | 29 (19.5%) | 43 (28.9%) |
| Borzoi | NHW | $\geq$ 0.9 | 117 | 31 (26.5%) | 26 (22.2%) | 24 (20.5%) | 36 (30.8%) |
| AlphaGenome | AA | $\geq$ 0.5 | 147 | 41 (27.9%) | 26 (17.7%) | 41 (27.9%) | 39 (26.5%) |
| AlphaGenome | AA | $\geq$ 0.6 | 131 | 35 (26.7%) | 25 (19.1%) | 37 (28.2%) | 34 (26.0%) |
| AlphaGenome | AA | $\geq$ 0.7 | 113 | 32 (28.3%) | 22 (19.5%) | 29 (25.7%) | 30 (26.5%) |
| AlphaGenome | AA | $\geq$ 0.8 | 100 | 28 (28.0%) | 19 (19.0%) | 25 (25.0%) | 28 (28.0%) |
| AlphaGenome | AA | $\geq$ 0.9 | 79 | 22 (27.8%) | 14 (17.7%) | 20 (25.3%) | 23 (29.1%) |
| AlphaGenome | CH | $\geq$ 0.5 | 113 | 15 (13.3%) | 28 (24.8%) | 28 (24.8%) | 42 (37.2%) |
| AlphaGenome | CH | $\geq$ 0.6 | 100 | 14 (14.0%) | 23 (23.0%) | 24 (24.0%) | 39 (39.0%) |
| AlphaGenome | CH | $\geq$ 0.7 | 82 | 8 (9.8%) | 20 (24.4%) | 24 (29.3%) | 30 (36.6%) |
| AlphaGenome | CH | $\geq$ 0.8 | 66 | 6 (9.1%) | 18 (27.3%) | 20 (30.3%) | 22 (33.3%) |
| AlphaGenome | CH | $\geq$ 0.9 | 51 | 4 (7.8%) | 14 (27.5%) | 15 (29.4%) | 18 (35.3%) |
| AlphaGenome | NHW | $\geq$ 0.5 | 336 | 77 (22.9%) | 83 (24.7%) | 71 (21.1%) | 105 (31.2%) |
| AlphaGenome | NHW | $\geq$ 0.6 | 255 | 58 (22.7%) | 60 (23.5%) | 53 (20.8%) | 84 (32.9%) |
| AlphaGenome | NHW | $\geq$ 0.7 | 200 | 50 (25.0%) | 46 (23.0%) | 39 (19.5%) | 65 (32.5%) |
| AlphaGenome | NHW | $\geq$ 0.8 | 165 | 43 (26.1%) | 38 (23.0%) | 30 (18.2%) | 54 (32.7%) |
| AlphaGenome | NHW | $\geq$ 0.9 | 130 | 33 (25.4%) | 28 (21.5%) | 25 (19.2%) | 44 (33.8%) |

Table S2: **Supplementary Table S2. Positive-class MAF distributions across ancestries and PIP thresholds.** For each model, ancestry, and PIP threshold, the table reports the number of scored positive variants, the median minor allele frequency (MAF), and the percentage of positives with MAF < 0.05 for the deduplicated manuscript positive benchmark. These summaries show that AA positives are consistently shifted toward lower MAF than CH and are generally comparable to or lower than NHW, depending on model and threshold.

| **Ancestry** | **PIP** | **Borzoi** | | | **AlphaGenome** | | |
|---|---|---|---|---|---|---|---|
| | | $n$ | **Median MAF** | **MAF < 0.05 (%)** | $n$ | **Median MAF** | **MAF < 0.05 (%)** |
| AA | $\geq 0.5$ | 140 | 0.229 | 12.1 | 147 | 0.228 | 12.9 |
| AA | $\geq 0.6$ | 124 | 0.235 | 12.1 | 131 | 0.234 | 13.0 |
| AA | $\geq 0.7$ | 106 | 0.239 | 14.2 | 113 | 0.237 | 15.0 |
| AA | $\geq 0.8$ | 93 | 0.232 | 16.1 | 100 | 0.229 | 17.0 |
| AA | $\geq 0.9$ | 72 | 0.220 | 16.7 | 79 | 0.214 | 17.7 |
| CH | $\geq 0.5$ | 113 | 0.268 | 4.4 | 113 | 0.268 | 4.4 |
| CH | $\geq 0.6$ | 100 | 0.268 | 5.0 | 100 | 0.268 | 5.0 |
| CH | $\geq 0.7$ | 82 | 0.261 | 6.1 | 82 | 0.261 | 6.1 |
| CH | $\geq 0.8$ | 66 | 0.251 | 7.6 | 66 | 0.251 | 7.6 |
| CH | $\geq 0.9$ | 51 | 0.239 | 7.8 | 51 | 0.239 | 7.8 |
| NHW | $\geq 0.5$ | 313 | 0.277 | 8.9 | 336 | 0.268 | 10.1 |
| NHW | $\geq 0.6$ | 236 | 0.266 | 9.3 | 255 | 0.260 | 11.0 |
| NHW | $\geq 0.7$ | 184 | 0.262 | 10.9 | 200 | 0.251 | 12.0 |
| NHW | $\geq 0.8$ | 149 | 0.223 | 13.4 | 165 | 0.221 | 14.5 |
| NHW | $\geq 0.9$ | 117 | 0.221 | 14.5 | 130 | 0.214 | 16.2 |

Table S3: **Supplementary Table S3. Summary of MAF-stratified, distance-matched AUROC across PIP thresholds.** Mean AUROC values are averaged across PIP thresholds 0.5–0.9 from the expanded 100× background analysis using available bootstrap estimates within each ancestry and MAF stratum. The $n$ range gives the minimum and maximum number of positive variants contributing across those thresholds. Rare-variant estimates, especially for CH ($n$ = 4–5), should be interpreted cautiously because of limited positive counts.

| **Ancestry** | **MAF bin** | **Borzoi** | | **AlphaGenome** | |
|---|---|---|---|---|---|
| | | **Mean AUROC** | $n$ **range** | **Mean AUROC** | $n$ **range** |
| AA | MAF < 0.05 | 0.790 | 12–17 | 0.675 | 14–19 |
| AA | $0.05 \leq$ MAF < 0.2 | 0.802 | 22–46 | 0.831 | 24–48 |
| AA | MAF $\geq 0.2$ | 0.833 | 38–77 | 0.816 | 41–80 |
| CH | MAF < 0.05 | 0.851 | 4–5 | 0.748 | 4–5 |
| CH | $0.05 \leq$ MAF < 0.2 | 0.596 | 12–24 | 0.694 | 12–24 |
| CH | MAF $\geq 0.2$ | 0.740 | 35–84 | 0.745 | 35–84 |
| NHW | MAF < 0.05 | 0.692 | 17–28 | 0.647 | 21–34 |
| NHW | $0.05 \leq$ MAF < 0.2 | 0.761 | 36–83 | 0.779 | 41–90 |
| NHW | MAF $\geq 0.2$ | 0.690 | 64–202 | 0.687 | 68–212 |

Table S4: **Supplementary Table S4. Sample sizes for FILER functional annotation overlap analysis.** Counts are deduplicated high-PIP positive variants available for each model, ancestry group, and PIP threshold.

| Model | PIP threshold | AA | CH | NHW |
|---|---|---|---|---|
| AlphaGenome | $\geq 0.5$ | 142 | 104 | 325 |
| AlphaGenome | $\geq 0.6$ | 126 | 93 | 245 |
| AlphaGenome | $\geq 0.7$ | 108 | 75 | 193 |
| AlphaGenome | $\geq 0.8$ | 96 | 60 | 158 |
| AlphaGenome | $\geq 0.9$ | 76 | 46 | 123 |
| Borzoi | $\geq 0.5$ | 136 | 104 | 309 |
| Borzoi | $\geq 0.6$ | 120 | 93 | 233 |
| Borzoi | $\geq 0.7$ | 102 | 75 | 183 |
| Borzoi | $\geq 0.8$ | 90 | 60 | 148 |
| Borzoi | $\geq 0.9$ | 70 | 46 | 116 |

Table S5: **Supplementary Table S5. Key FILER functional annotation overlap among high-PIP variants.** Values are percentages of high-PIP positive variants overlapping selected FILER categories emphasized in the manuscript. Chi-square $q$ values are Benjamini–Hochberg FDR-adjusted 3×2 tests comparing AA, CH, and NHW within each model and PIP threshold. For readability, this compact table reports the least and most stringent PIP thresholds.

| **Model** | **PIP** | **Category** | **AA (%)** | **CH (%)** | **NHW (%)** | **Chi-square $q$** |
|---|---|---|---|---|---|---|
| AlphaGenome | $\geq 0.5$ | Accessible chromatin | 70.4 | 63.5 | 54.8 | 0.021 |
| AlphaGenome | $\geq 0.5$ | Hi-C | 76.8 | 74.0 | 63.1 | 0.021 |
| AlphaGenome | $\geq 0.5$ | IM-PET | 48.6 | 43.3 | 34.5 | 0.034 |
| AlphaGenome | $\geq 0.5$ | pcHi-C | 90.1 | 90.4 | 85.8 | 0.327 |
| AlphaGenome | $\geq 0.5$ | eQTL | 83.1 | 96.2 | 95.1 | 1.86e-04 |
| AlphaGenome | $\geq 0.5$ | sQTL | 48.6 | 73.1 | 62.5 | 0.003 |
| AlphaGenome | $\geq 0.9$ | Accessible chromatin | 78.9 | 67.4 | 65.9 | 0.393 |
| AlphaGenome | $\geq 0.9$ | Hi-C | 76.3 | 65.2 | 59.3 | 0.186 |
| AlphaGenome | $\geq 0.9$ | IM-PET | 52.6 | 47.8 | 43.1 | 0.573 |
| AlphaGenome | $\geq 0.9$ | pcHi-C | 92.1 | 84.8 | 78.0 | 0.163 |
| AlphaGenome | $\geq 0.9$ | eQTL | 81.6 | 97.8 | 93.5 | 0.053 |
| AlphaGenome | $\geq 0.9$ | sQTL | 50.0 | 65.2 | 60.2 | 0.429 |
| Borzoi | $\geq 0.5$ | Accessible chromatin | 69.9 | 63.5 | 55.7 | 0.056 |
| Borzoi | $\geq 0.5$ | Hi-C | 77.2 | 74.0 | 63.4 | 0.034 |
| Borzoi | $\geq 0.5$ | IM-PET | 49.3 | 43.3 | 35.6 | 0.062 |
| Borzoi | $\geq 0.5$ | pcHi-C | 90.4 | 90.4 | 87.7 | 0.647 |
| Borzoi | $\geq 0.5$ | eQTL | 84.6 | 96.2 | 96.8 | 5.75e-05 |
| Borzoi | $\geq 0.5$ | sQTL | 49.3 | 73.1 | 63.1 | 0.005 |
| Borzoi | $\geq 0.9$ | Accessible chromatin | 78.6 | 67.4 | 69.0 | 0.468 |
| Borzoi | $\geq 0.9$ | Hi-C | 77.1 | 65.2 | 60.3 | 0.242 |
| Borzoi | $\geq 0.9$ | IM-PET | 54.3 | 47.8 | 45.7 | 0.709 |
| Borzoi | $\geq 0.9$ | pcHi-C | 92.9 | 84.8 | 80.2 | 0.242 |
| Borzoi | $\geq 0.9$ | eQTL | 84.3 | 97.8 | 95.7 | 0.075 |
| Borzoi | $\geq 0.9$ | sQTL | 51.4 | 65.2 | 61.2 | 0.468 |

Table S6: **Supplementary Table S6. Selected pairwise ancestry comparisons for key FILER annotation categories.** Rows show pairwise Fisher exact tests with FDR-adjusted $q < 0.10$ for the compact set of categories discussed in the manuscript at PIP thresholds shown in Supplementary Table S5. Fractions are percentages of high-PIP variants overlapping each category; difference is ancestry 1 minus ancestry 2 in percentage points.

| **Model** | **PIP** | **Category** | **Comparison** | **Frac. 1 (%)** | **Frac. 2 (%)** | **Diff.** | **$q$** |
|---|---|---|---|---|---|---|---|
| AlphaGenome | $\geq 0.5$ | Accessible chromatin | AA vs NHW | 70.4 | 54.8 | +15.7 | 0.017 |
| Borzoi | $\geq 0.5$ | Accessible chromatin | AA vs NHW | 69.9 | 55.7 | +14.2 | 0.050 |
| AlphaGenome | $\geq 0.5$ | Hi-C | AA vs NHW | 76.8 | 63.1 | +13.7 | 0.034 |
| Borzoi | $\geq 0.5$ | Hi-C | AA vs NHW | 77.2 | 63.4 | +13.8 | 0.050 |
| AlphaGenome | $\geq 0.5$ | IM-PET | AA vs NHW | 48.6 | 34.5 | +14.1 | 0.038 |
| Borzoi | $\geq 0.5$ | IM-PET | AA vs NHW | 49.3 | 35.6 | +13.7 | 0.054 |
| AlphaGenome | $\geq 0.5$ | eQTL | AA vs CH | 83.1 | 96.2 | -13.1 | 0.016 |
| AlphaGenome | $\geq 0.5$ | eQTL | AA vs NHW | 83.1 | 95.1 | -12.0 | 0.002 |
| Borzoi | $\geq 0.5$ | eQTL | AA vs CH | 84.6 | 96.2 | -11.6 | 0.050 |
| Borzoi | $\geq 0.5$ | eQTL | AA vs NHW | 84.6 | 96.8 | -12.2 | 8.17e-04 |
| AlphaGenome | $\geq 0.5$ | sQTL | AA vs CH | 48.6 | 73.1 | -24.5 | 0.003 |
| AlphaGenome | $\geq 0.5$ | sQTL | AA vs NHW | 48.6 | 62.5 | -13.9 | 0.038 |
| Borzoi | $\geq 0.5$ | sQTL | AA vs CH | 49.3 | 73.1 | -23.8 | 0.005 |
| Borzoi | $\geq 0.5$ | sQTL | AA vs NHW | 49.3 | 63.1 | -13.8 | 0.050 |
